# Fibroblast activation protein enzyme deficiency prevents liver steatosis, insulin resistance and glucose intolerance and increases fibroblast growth factor-21 in diet induced obese mice

**DOI:** 10.1101/460279

**Authors:** Sumaiya Chowdhury, Sunmi Song, Hui Emma Zhang, Xin Maggie Wang, Margaret G. Gall, Denise Ming Tse Yu, Angelina J. Lay, Michelle Sui Wen Xiang, Kathryn A. Evans, Stefanie Wetzel, Yolanda Liu, Belinda Yau, Andrew L. Coppage, Lisa Lo, Rebecca A. Stokes, Wayne J. Hawthorne, Gregory J. Cooney, Susan V. McLennan, Jenny E. Gunton, William W. Bachovchin, Nigel Turner, Melkam A. Kebede, Geoffrey W. McCaughan, Stephen M. Twigg, Mark D. Gorrell

## Abstract

**Background & Aims:** Fibroblast activation protein-a (FAP) is a post-proline peptidase closely related to dipeptidyl peptidase-4. FAP degrades bioactive peptides including fibroblast growth factor-21 (FGF-21) and neuropeptide Y. We examined metabolic outcomes of specific genetic ablation of FAP and its enzyme activity in a mouse model of diet-induced obesity (DIO) causing fatty liver.

**Methods:** Wildtype (WT) and genetically modified FAP deficient mice that specifically lacked either the FAP protein or FAP enzyme activity received chow, or an atherogenic diet for 8 to 20 weeks of DIO.

**Results:** FAP deficient male and female mice in the DIO model were more metabolically healthy than controls. The FAP deficient mice had less glucose intolerance, liver lipid, adiposity, insulin resistance, pancreatic and plasma insulin, pancreatic β-cell hyperplasia, serum alanine transaminase and circulating cholesterol compared to wild type controls. Furthermore, FAP deficiency lowered respiratory exchange ratio and greatly increased intrahepatic non-esterified free fatty acids, indicative of increased lipolysis and β-oxidation. Concordantly, lipogenic genes (*Ppar*g, *Gck, Acc, Fasn*) and hepatic triglyceride and fatty acid uptake genes (*Cd36*, *Apoc3, Ldlr*) and plasma low-density lipoprotein cholesterol were downregulated. Glucagon like peptide-1 levels were unaltered. FAP was localized to human pancreatic β-cells and pancreas from diabetes mellitus patients contained elevated FAP activity. Comparable data from a FAP gene knockout mouse and a novel mouse lacking FAP enzyme activity indicated that these metabolic changes depended upon the enzymatic activity of FAP. These changes may be driven by FGF-21, which was upregulated in livers of FAP deficient DIO mice.

**Conclusion:** This is the first study to show that specific genetic ablation of FAP activity or protein protects against DIO-driven glucose intolerance, hyperinsulinaemia, insulin resistance, hypercholesterolaemia and liver steatosis in mice and provide mechanistic insights.

## Introduction

Fibroblast Activation Protein-alpha (FAP) is a unique extracellular serine protease of the dipeptidyl peptidase-4 (DPP4) family. FAP degrades many bioactive peptides [1]. Neuropeptide Y (NPY) and fibroblast growth factor-21 (FGF-21) are physiological substrates [1-6]. FAP expression is very low in normal adult tissues [7, 8] but elevated in cirrhosis [8, 9] and non-alcoholic steatohepatitis (NASH) [10], and greatly upregulated in activated fibroblasts at sites of tissue remodeling [9]. FAP may be pro-fibrotic in lung [11]. FAP has no fundamental effect on the immune system [12].

DPP4, the closest relative of FAP, is the best-characterised member of the DPP4 family. DPP4 inhibitors are a successful therapy for type 2 diabetes mellitus [13]. Depleting DPP4 enzyme activity decreases glucose intolerance, insulin resistance, obesity and liver steatosis in mice [14-18], mainly due to increased active glucagon like peptide-1 (GLP-1) and altered lipid metabolism.

The role of FAP in energy metabolism is unexplored. A recent study found that non-selective FAP inhibition using Val-boro-Pro improved glucose tolerance and insulin sensitivity, lowered body weight, food intake, adiposity and cholesterol and increased energy expenditure [6]. Because Val-boro-Pro is non-selective, those effects could be due to inhibition of other DPPs including DPP4, and so investigations using FAP-selective approaches are needed. FAP cleaves several hormones [3, 19] and rapidly inactivates FGF-21 in human and increases FGF-21 degradation in mouse [1, 5, 6, 20]. FGF-21 improves glucose and insulin homeostasis and energy metabolism in non-alcoholic fatty liver disease (NAFLD).

With specific molecular genetic deficiency of FAP in two mouse strains and a model of diet-induced obesity (DIO), insulin resistance and liver steatosis, we found that FAP deficiency lessens glucose intolerance, insulin resistance and steatosis. Our mechanistic investigations indicated that FAP deficiency acts differently to DPP4, by increasing fat burning and with increased *Fgf-21* and *Lipin-1* expression, and increased FGF-21, and greater ACC phosphorylation.

## METHODS

### Ethics

Studies involving human islets accorded with approval LNR/15/WMEAD/386 from the Western Sydney Health District Ethics Committee. Studying de-identified human pancreas samples from the nPOD tissue consortium was approved by the Tufts University Health Science Campus Institutional Review Board.

All animal handling and experimental procedures were approved by the Animal Ethics Committees of the University of Sydney and the Sydney Local Health District (SLHD) under protocol 2013-017 or the Garvan Institute/St Vincent’s Hospital Animal Experimentation Ethics Committee under protocol 11/41. Mice in experiments were housed in animal facilities of SLHD, Centenary Institute or Garvan Institute in compliance with the Australian Code of Practice for the Care and Use of Animals for Scientific Purpose. Wild type (WT) C57BL/6J mice were obtained from either the Animal Resource Center (ARC; Perth, WA, Australia) or Australian BioResources (Moss Vale, NSW, Australia).

The FAP gene knockout (gko) mouse strain, which lacks FAP protein [21] (gift from Boehringer Ingelheim, Germany), were backcrossed twice onto C57BL/6J for more than seven generations [12]. The new FAP gene knock-in (gki) mouse strain was generated as described in Fig.S1. All ES clones and mice were screened and verified by Southern Blot analyses (Fig. S2A-D). FAPgki mice lacked FAP enzyme activity (Fig. S2E) but retained cell surface expression of FAP protein (Fig. S3). Age-matched adult male and female mice were housed at 21°C under a 12 h light 12 h dark cycle, offered water and food *ad libitum*, as described [22, 23]. HFD was purchased (Cat. No. SF03-020; Specialty Feeds) unless specified as in-house HFD (see Supplementary Methods).

### Glucose and insulin tolerance tests

For glucose tolerance test (GTT) and insulin tolerance test (ITT), mice were fasted for 6 hours (8am-2pm) [24]. GTT used D-glucose at 2 g/kg (Gibco TM, Auckland, NZ) administered by either oral gavage or intraperitoneal (ip) as stated. ITT used ip insulin (recombinant NovoRapid^®^ insulin; 0.75 U/kg, Novo Nordisk, New Zealand, or 0.5 U/kg human insulin, Sigma-Aldrich CasNo.11061-68-0). Glucose was measured in tail vein blood using an Accu-Chek Glucometer (Roche, Diagnostics GmbH, Mannheim, Germany).

### Organ collection

Mice were euthanized by carbon dioxide inhalation. Blood was collected by cardiac puncture in anti-coagulant free 1 mL Z-serum capillary blood collection tubes (Catalogue No.450474; Greiner Bio-One GmbH, Kremsmunster, Austria). After 30-60 min for clotting, blood was centrifuged for 5 min at 3000 x g for −80°C serum storage. Liver, abdominal white adipose tissue (WAT) and interscapular brown adipose tissue (BAT) were weighed and samples were snap frozen or formalin-fixed. Identical liver lobes were sampled from each mouse for each procedure.

### Blood biochemistry and hormone analyses

Liver function tests, which included alanine transaminase (ALT), aspartate aminotransferase (AST) and circulating cholesterol, were performed on serum samples by the Clinical Biochemistry Department of RPAH. Serum insulin was measured using a rat/mouse insulin ELISA kit according to manufacturer’s instructions (CatNo.EZRMI-13K; Merck Millipore, Darmstadt, Germany). Circulating adiponectin was measured by ELISA kit (Catalogue No. 80569; Crystal Chem, IL, USA). Spectrophotometric measurements used a POLARstar Omega plate reader (BMG Labtech, Offenburg, Germany). Insulin resistance was estimated according to the Homeostasis Model Assessment (HOMA-IR). The calculations were done with fasting blood glucose (mmol/l) and serum insulin (pmol/l) using the HOMA calculator (www.dtu.ox.ac.uk/homacalculator/ The University of Oxford; Accessed 30^th^ October 2014).

### Histology and immunohistochemistry

Tissue embedding and Haematoxylin-Eosin (H&E) and Sirius Red staining [22, 25] were performed by the Histopathology Department, University of Sydney. Immunohistochemistry procedures on mouse tissues have been described previously [23, 25, 26].

### Human islet recovery, dispersion and immunocytochemistry

Human islets isolated as described [27] were suspended in RPMI1640 media, 11 mM glucose, 10% fetal bovine serum for overnight incubation then dispersed in TrypLE Express for 3 min, briefly dissociated into single cells by pipetting, resuspended into 5 volumes of supplemented RPMI1640, then placed on glass coverslips. After overnight incubation, cells were washed twice in Wash Buffer (0.1% BSA in PBS with 0.01% sodium azide), fixed at ambient temperature for 20 min with 4% paraformaldehyde in PBS, then washed twice in Wash Buffer before antigen retrieval in 0.1% SDS for 5 min. Cells were washed, then blocked for 1h with Serum-Free Protein Block (DAKO, X0909) before overnight staining with guinea pig anti-insulin (DAKO, A0564) and mouse anti-glucagon (Sigma, SAB4200685), 3 washes, staining with fluorescently-labelled anti-guinea pig and anti-mouse secondary antibodies, and mounting onto microscope slides with Prolong Diamond Antifade Mountant containing DAPI (Thermo Fisher, P36965). Slides were imaged using a Leica SP8 confocal microscope.

### Semi-quantitative oil red O (ORO) assay

Total liver lipid was measured by ORO semi-quantitative assay, as previously described [22]. The ORO stock solution 0.25% (wt/vol) was freshly diluted in 10% dextran at 6:4 (ORO: dextran) then 30-50 mg (ww) liver was homogenized and incubated in the ORO working solution for 1 h. After washing with 60% isopropanol to remove excess dye, the dye incorporated into lipid was extracted in 99% isopropanol. Absorbance at 520 nm was measured alongside an ORO standard curve.

### Non-esterified free fatty acid (NEFA) measurement

The NEFA C kit (Wako Diagnostics, Osaka, Japan) was used as instructed by the manufacturer. Liver lipid for the NEFA assay was extracted from snap frozen liver tissue in isopropanol (50 mg ww/mL).

### Real-time quantitative polymerase chain reaction (qPCR)

RNA extraction used TRIzol (Cat no. 15596026; Invitrogen), 0.5 μg RNA was reverse transcribed to cDNA using Superscript^®^ VILO^TM^ cDNA synthesis kit (Cat no. 11754; Invitrogen), real-time qPCR with Taqman^®^ gene expression assays (Applied Biosystems; Table S1) was performed using the Stratagene^®^ Mx3000P™ System (La Jolla, CA, USA) and gene expression quantified using standard curves, as described previously [12, 23, 28].

### Indirect calorimetry

Indirect calorimetry was applied to age-matched female WT and FAPgko mice on chow for 20 weeks or in-house HFD [23] for 12 weeks. Whole-body oxygen consumption rate (VO_2_) and respiratory exchange ratio (RER) of individual mice were measured using an eight-chambered indirect calorimeter (Oxymax series; Columbus Instruments, Columbus, OH) as previously described [29].

### FAP enzyme activity assay

Fresh frozen mouse tissue and human pancreas samples from the nPOD Consortium tissue bank were homogenized in ice-cold lysis buffer. Human pancreas in lysis buffer (50 mM Tris-HCL pH 7.5, 10% glycerol, 5 mM EDTA, 1 mM DTT) was homogenized at 10 mL/g, and assayed at 1 mg protein/mL with 100 μM 3144-AMC [30]. Otherwise, the lysis buffer was 50 mM Tris-HCl pH 7.6, 1 mM EDTA, 10% glycerol, 1% Triton-X114, with complete protease inhibitors (Roche, Basel, Switzerland) and the soluble supernatant retained for FAP enzyme activity assay, performed using 100 μg (protein) of tissue lysate and fluorogenic substrate 3144-AMC at 150 μM, as described previously [7].

### Immunoblotting

The 3T3-L1 cells were differentiated into adipocytes as described [31]. These adipocytes were incubated with full-length or FAP-cleaved FGF-21 at 500 ng/mL. FGF-21 was pre-incubated with PBS or recombinant FAP (substrate: enzyme ratio = 5:1) for 2 h, with or without FAP-selective inhibitor ARI-3099 [32] at 5 µM. Immunoblots of FGF-21, Erk, phosphorylated Erk (p-Erk) ACC, pACC, PGC-1α and β-actin were performed on lysates of tissues or of cells.

### Faecal lipid

Faecal pellets were collected from mouse bedding (n = 3 cages per group) at 19 weeks of HFD and faecal fat quantified as described [33]. 0.5 g of faeces was dissolved in 2 mL deionized water overnight at 4°C then homogenized by vortexing, then lipid extraction was performed with methanol-chloroform (2:1 v/v). The extracted lipophilic layer (lower chloroform phase) was vacuum dried at 45°C for 2 h. The dry pellet (lipid) was weighed to calculate faecal lipid as mg lipid per g faeces.

### Statistical analysis

Statistical analyses in GraphPad Prism (v.7) used non-parametric Mann-Whitney U test and significance was assigned to p-values less than 0.05.

## RESULTS

### FAPgko mice were protected against HFD induced impaired glucose homeostasis and insulin resistance

On chow diet, FAPgko and WT mice had similar glucose tolerance (Fig. 1A). In contrast, from 8 weeks of HFD, FAPgko mice were resistant to the impaired glucose tolerance seen in WT mice (Fig. 1B-D). These female FAPgko mice were able to decrease blood glucose faster than WT mice from 30 minutes after oral glucose challenge (Fig. 1B-D). Concordantly, FAPgki mice on HFD for 12 weeks also had improved glucose tolerance compared to WT HFD controls (Fig. 1E-F). This improved glucose tolerance was seen in both male and female FAP deficient mice (Fig. S4, Fig. 1).

**Fig. 1.**
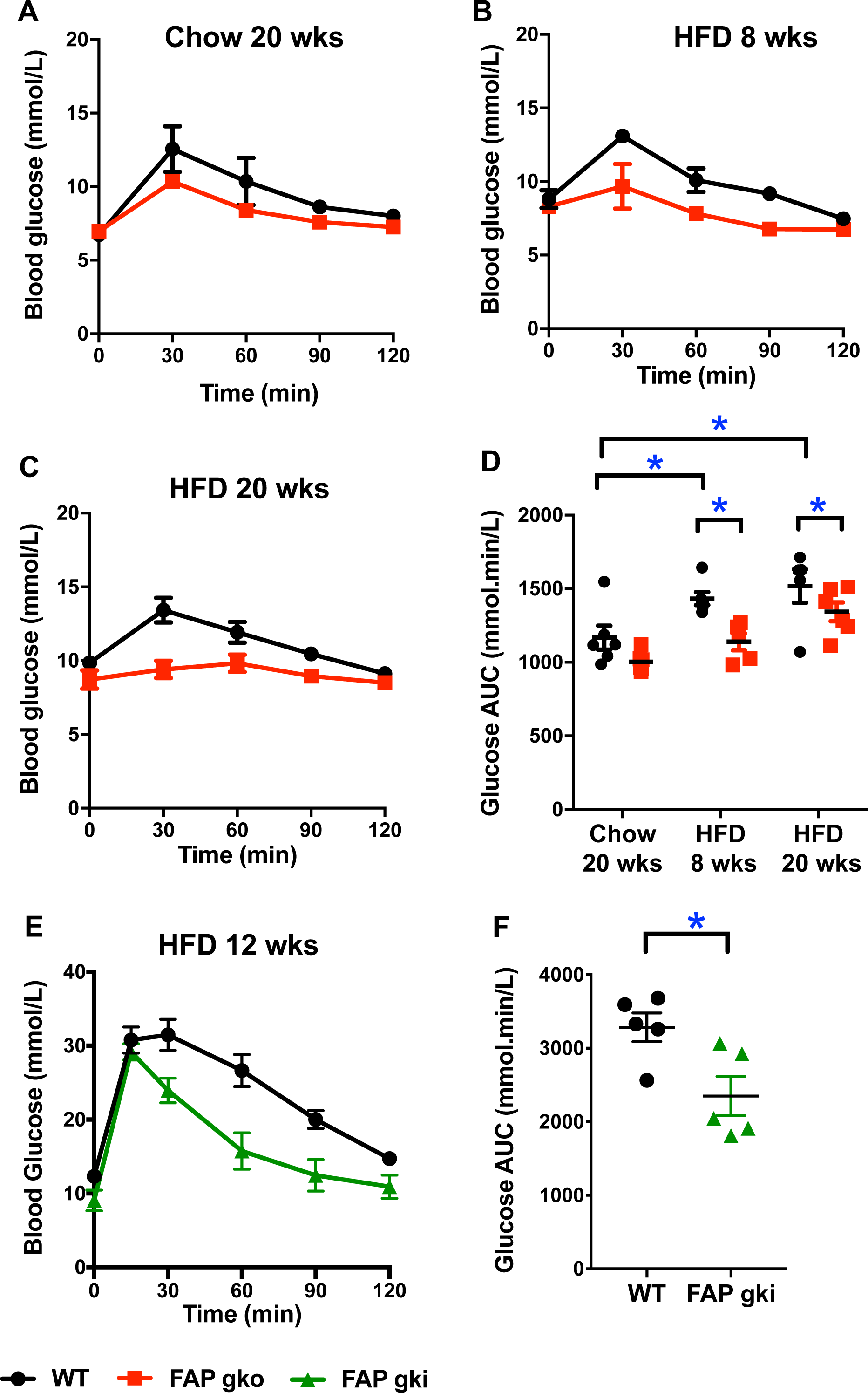
Improved glucose tolerance in FAP deficient DIO mice. Blood glucose following oral (A-D) or intraperitoneal (E-F) GTT at 20 weeks of chow (A), 8 weeks of HFD (B), 20 weeks of HFD (C), 12 weeks of in-house HFD (E), and glucose AUC (D, F), in female FAPgko (A-D) or male FAPgki (E-F) mice compared to WT mice. Individual replicates (D, F). Mean ± SEM (A-F). n=5-6 mice per group. *p<0.05 versus genotype-matched controls using Mann-Whitney U test.

On chow diet, FAPgko mice had insulin tolerance levels similar to WT mice (Fig. 2A and 2D). However, at 20 weeks of HFD, insulin manifested significantly greater glucose lowering efficacy in FAPgko mice than WT mice (Fig. 2B-D). FAPgki mice on 16 weeks of HFD had a similarly improved insulin tolerance (Fig. 2E-F). These data suggest that obesity-linked insulin resistance in mice requires FAP enzyme activity.

**Fig. 2.**
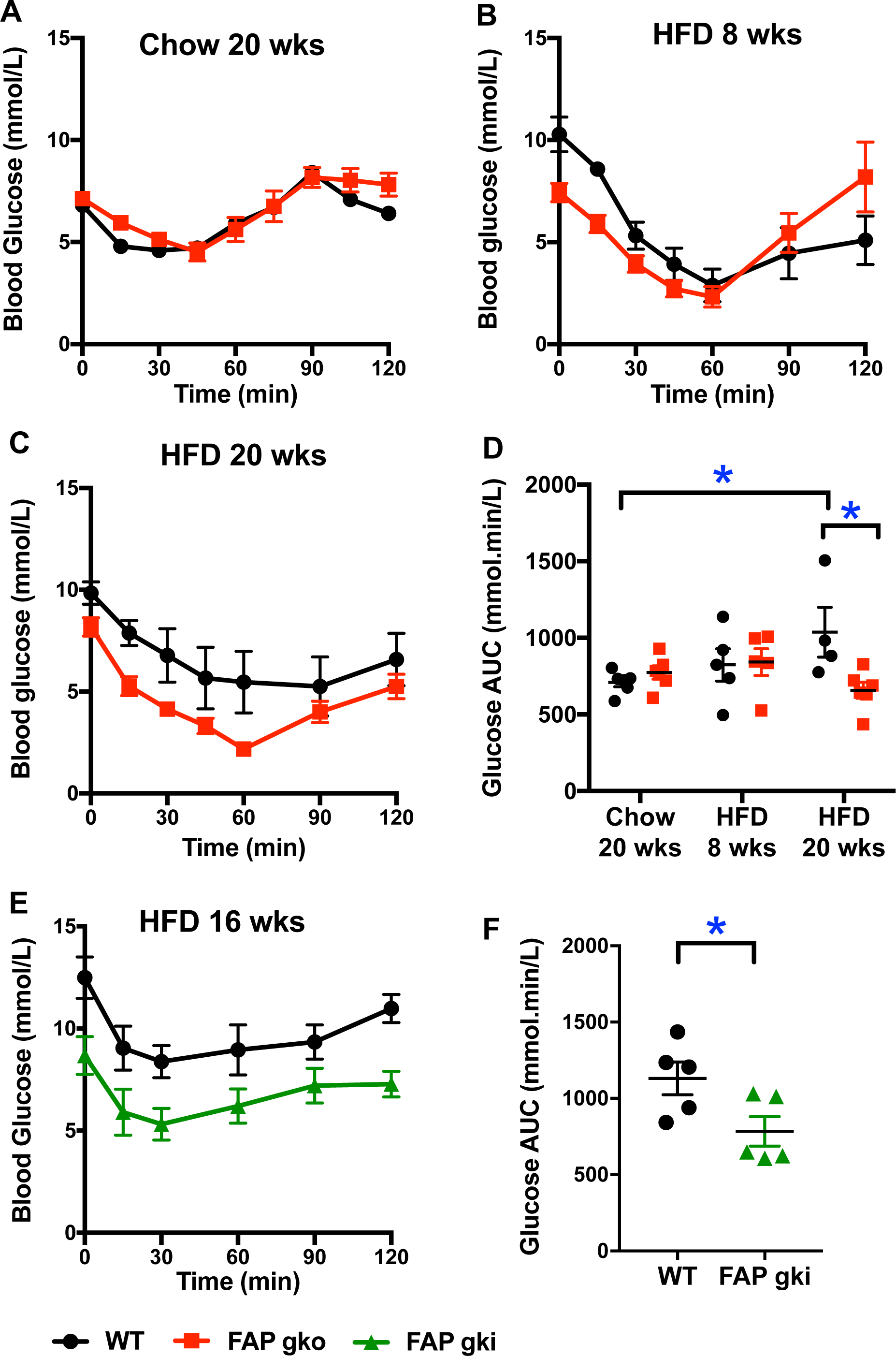
Insulin sensitivity in FAP deficient DIO mice. Insulin tolerance test (ITT), showing blood glucose following an intraperitoneal ITT at 20 weeks of chow (A), 8 weeks of HFD (B), 20 weeks of HFD (C), 16 weeks of in-house HFD (E) and glucose AUC (D, F) in female FAPgko (A-D) or male FAPgki (EF) mice, and WT controls. Individual replicates and mean ± SEM. n= 5-6 mice per group. *p<0.05 by Mann-Whitney U test.

WT HFD mice exhibited a 3-fold increase in random plasma insulin level at 8 and 20 weeks of HFD compared to WT chow mice, indicative of hyperinsulinemia (Fig. 3A-B). Both FAPgko chow and FAPgko HFD mice maintained insulin levels similar to the chow fed WT mice, showing that FAPgko mice were resistant to HFD induced hyperinsulinemia (Fig. 3A-B). Moreover, FAPgki HFD mice resisted hyperinsulinemia (Fig. 3C) and, concordantly, exhibited less pancreatic β-cell hypertrophy compared to WT (Fig. 3D; Fig. S5).

**Fig. 3.**
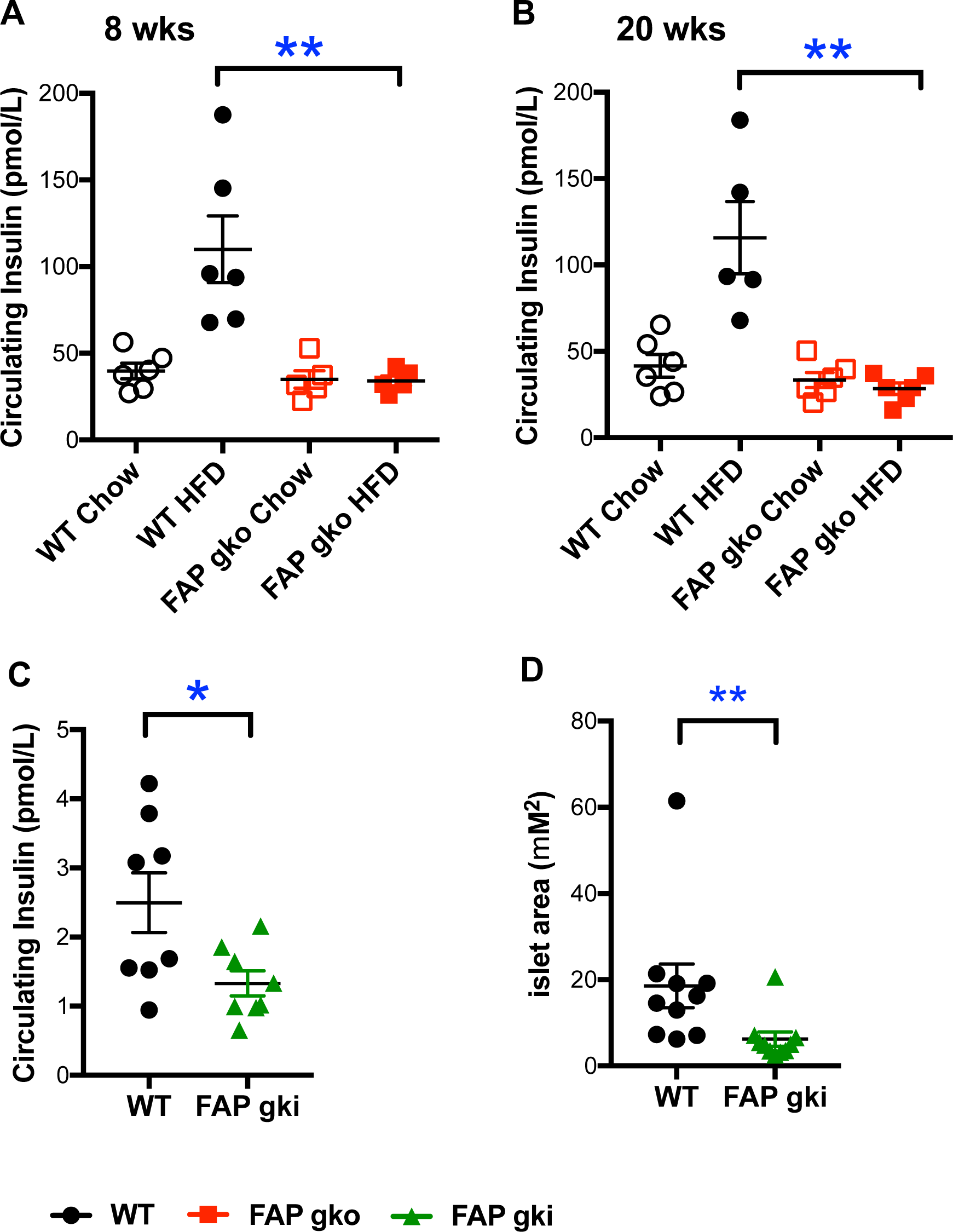
FAP deficient DIO mice were protected against insulin resistance. FAPgko (A-B) and FAPgki (C-D) mice following 8 weeks (A) or 20 weeks (B) of HFD or 12 weeks of in-house HFD (C-D) did not have elevated circulating insulin (A-C), and had less β-cell hypertrophy (D). Basal insulin level was measured in overnight - fasted female (A-B) or male (C-D) mice. Individual replicates and mean ± SEM. n=5-6 mice per group. *p<0.05, **p<0.01 by Mann-Whitney U test.

HOMA was used as a surrogate method for estimating insulin resistance. The relationship between glucose and insulin in the basal state reflects the balance between hepatic glucose output and insulin secretion. HOMA-IR scores were 0.84±0.13 and 0.68±0.09 on chow and 2.38±0.37 and 0.58±0.07 on HFD from WT and FAPgko mice, respectively. Thus, FAPgko mice on either diet had similar HOMA IR values as WT chow mice, indicating that FAPgko mice were protected against the obesity linked insulin resistance that occurred in the WT HFD mice. Similarly, HOMA-IR was 1.23±0.21 in WT and 0.62±0.33 in FAPgki mice on HFD. Therefore, HFD-induced insulin resistance, observed in WT mice, requires FAP enzyme activity.

### FAPgko mice were protected from HFD induced liver steatosis

Chow fed WT and FAPgko mice had similar liver histology, healthy hepatocytes and normal liver architecture (Fig. 4A-B). Eight weeks of HFD promoted onset of liver steatosis, as evidenced by the presence of intrahepatic vacuoles (clear areas) (Fig. 4C; arrows) and elevated Oil-Red O stain (Fig. S6) in WT mice. Macro and microvesicular steatosis occurred in WT mice, however FAPgko mice had fewer macrovesicular and very little microvesicular steatosis compared to WT HFD mice (Fig. 4D). The macrovesicular steatosis in FAPgko mice was mainly localized near the portal tracts, whereas in the central zone surrounding the central vein hepatocytes and other histological features were normal.

**Fig. 4.**
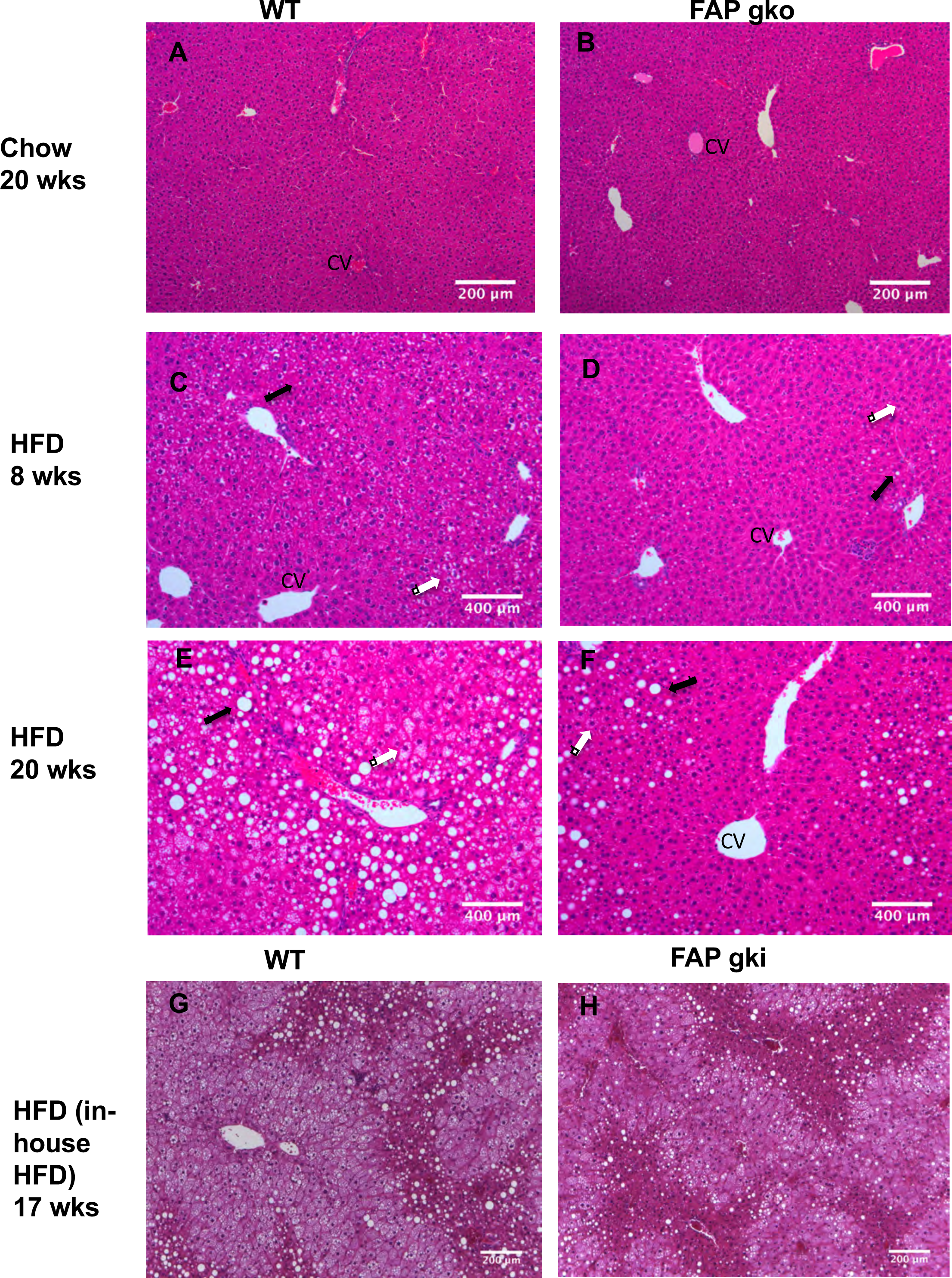
FAP deficient DIO mice were protected from HFD induced liver steatosis. Representative liver sections (H&E) from WT (A, C, E, G), FAPgko (B, D, F) and FAPgki (H) mice at 20 weeks of chow (A-B), 8 weeks of HFD (CD), 20 weeks of HFD (E-F), or 17 weeks of in-house HFD (G-H). Arrows indicate macrovesicular (black) and microvesicular (white) steatosis. CV=central vein. Scale bars=200 µm.

At 20 weeks of HFD, WT mice exhibited extensive steatosis with hepatocyte ballooning accompanied by reduced hepatic sinusoidal spaces (Fig. 4E; arrows). In contrast, FAPgko mice had significantly less steatosis and hepatocyte ballooning (Fig. 4F). The macrovesicular steatosis was mainly localized near the portal tracts, with healthy hepatocytes and normal histology surrounding the central vein. Similarly, male FAPgki mice had less liver steatosis compared to WT mice on HFD (Fig. 4G-H). Oil-Red O analysis showed less lipid in FAPgko liver, compared to WT liver, after 20 weeks of HFD (Fig. S6). These results show that FAPgko mice were protected from HFD induced liver steatosis.

HFD did not induce liver inflammation or fibrosis in either WT or FAPgko mice, as evident from both H&E and Sirius red stained sections (Fig. 4C-F; Fig. S7). Measures of hepatic damage, ALT and AST, were elevated in HFD fed mice (Fig. 5A,B). However, FAPgko mice exhibited lesser increases in ALT and AST compared with WT mice, suggesting that the FAPgko mice were protected against HFD induced liver injury. Tissue macrophage densities were low in liver, BAT and WAT, but greater in FAPgki than WT liver (Fig. S8).

**Fig. 5.**
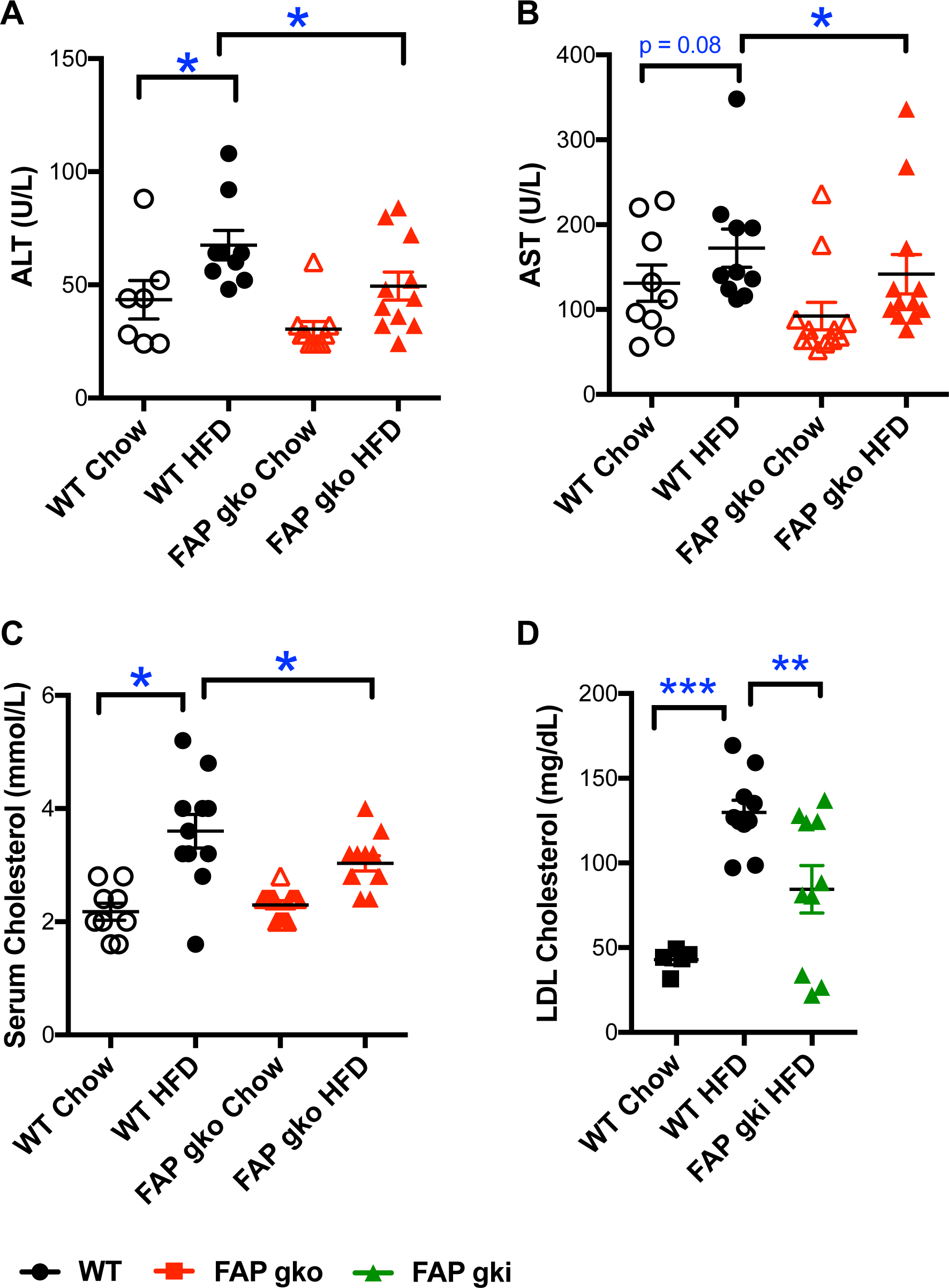
FAP deficient DIO mice were protected from HFD induced liver injury. ALT (A), AST (B), total cholesterol (C), LDL cholesterol (D) from WT and FAPgko (A-C) and FAPgki (D) overnight-fasted mice at 20 weeks of HFD (A-C) or 12 weeks of in-house HFD (D), and chow controls. Individual replicates and mean ± SEM. n=10-12. *p<0.05, **p<0.01 by Mann-Whitney U test.

Increased circulating total cholesterol exacerbates liver steatosis and inflammation and is a crucial driver towards NASH. FAPgko and FAPgki mice on HFD had lower circulating total cholesterol (Fig. 5C) and LDL cholesterol (Fig. 5D) compared to WT mice.

### FAP deficiency lessened adiposity

FAPgko mice and WT mice had similar body weights on chow. FAPgko mice on HFD had mildly less body weight and body weight gain compared to WT (Fig. S9A, B). FAPgki mice on in-house HFD had less body weight but similar rate of weight gain as WT mice (Fig. S9C, D). Both FAPgko and FAPgki HFD mice had less white adipose tissue (WAT) compared to WT HFD mice (Fig. 6A-E). Among the three abdominal fat depots collected for WAT, the most significant difference was seen in the gonadal WAT. FAPgki HFD mice had 30% less gonadal WAT than WT HFD mice (p<0.005) (Fig. 6E), and had elevated plasma adiponectin (Fig. 6G,H).

**Fig. 6.**
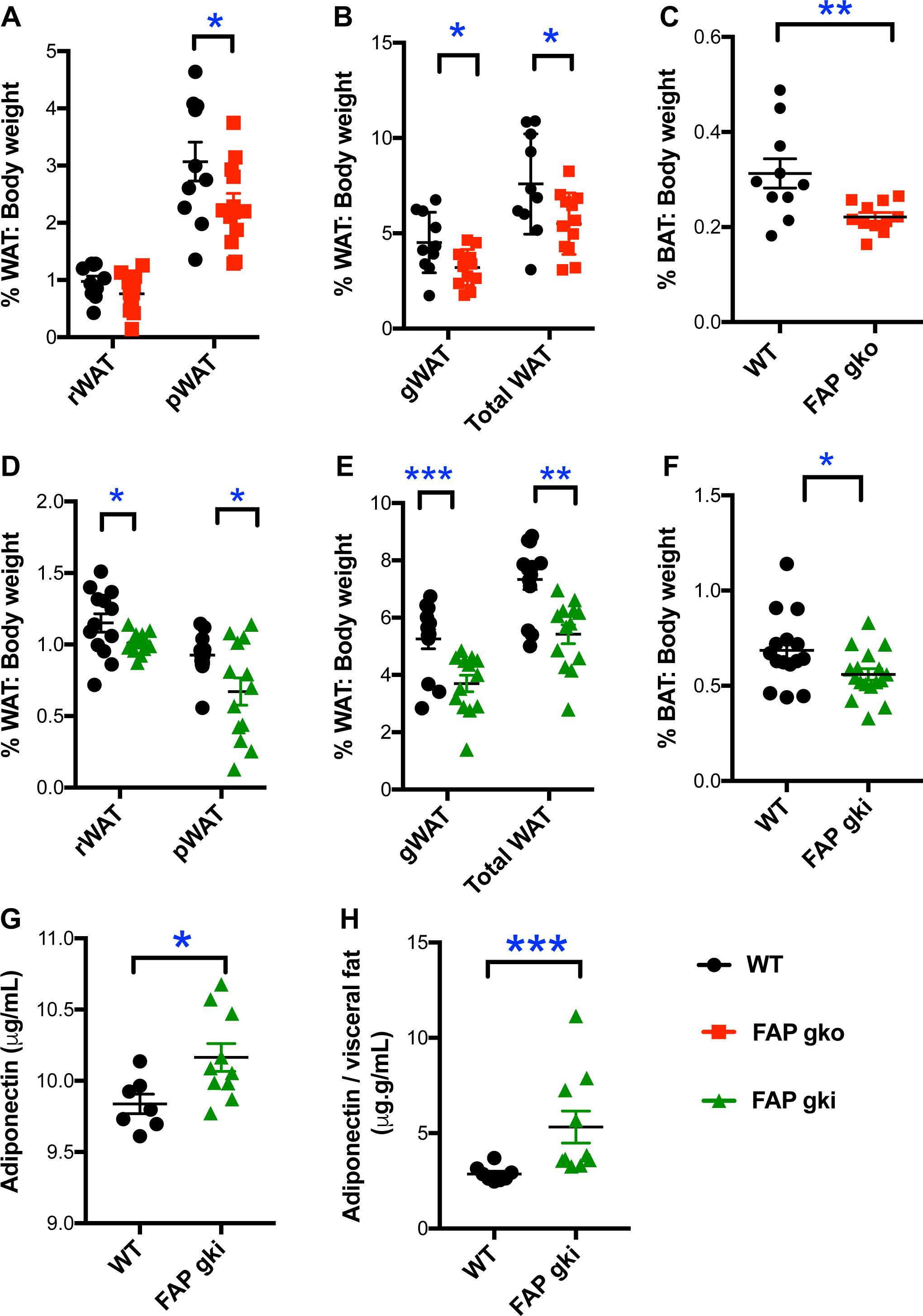
Less adiposity in FAP deficient DIO mice. Compared with WT, FAPgko mice (A-C) and FAPgki mice (D-H) had less abdominal white adipose tissue (WAT) and interscapular brown adipose tissue (BAT) as a percentage of body weight. rWAT = retroperitoneal WAT, pWAT = perirenal WAT, gWAT = gonadal WAT. Serum adiponectin concentration (G) and normalized to visceral fat weight (H) in WT and FAPgki mice at 12 weeks of in-house HFD. Individual replicates and mean ± SEM. 20 weeks of HFD, n = 10-12 females (A-C), or 17 weeks of in-house HFD, n=12-14 males (D-H). *p<0.05, **p<0.01, ***p<0.001.

FAPgko and FAPgki HFD mice had less brown adipose tissue (BAT) compared to WT HFD mice (Fig. 6C,F). FAP enzyme activity was greater in BAT from WT HFD mice compared to WT chow mice (Fig. 7A). There was no difference in FAP activity in WAT and pancreas in the WT chow versus WT HFD mice. WT HFD mouse liver had less FAP activity compared to WT chow liver (Fig. 7).

**Fig. 7.**
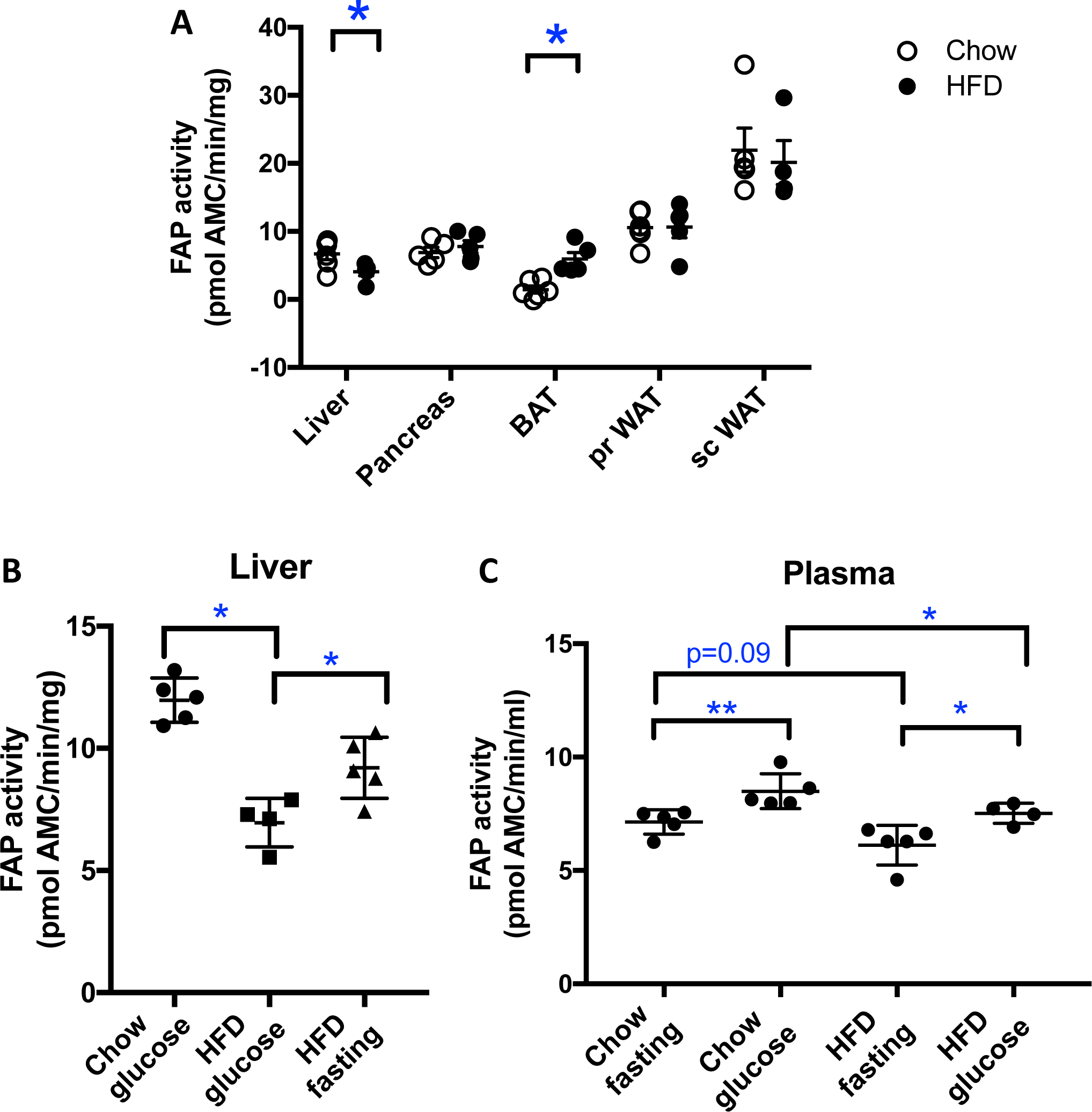
FAP activity in WT mice at 20 weeks of diet. Tissue (A, B) and plasma (C) samples from 5-6 female WT mice per group. Individual replicates and mean ± SEM. *p<0.05

Weight matched FAPgko mice had improved glucose tolerance compared to WT mice, on the in-house HFD (Fig. S9A-D). Moreover, regression analysis of GTT glucose AUC against body weight and adiposity showed that FAPgko and FAPgki mice had less glucose AUC than WT mice of comparable body weights and adiposity on either HFD (Fig. S10). With or without a body weight difference, FAPgko mice had improved liver histology compared to WT mice at 20 weeks of in-house HFD (Fig. 4A,B; Fig. S7A, B). Thus, the role of FAP in glucose homeostasis and liver steatosis in diet-induced obesity is independent of body weight gain or adiposity.

The major mechanism of metabolic improvement in DPP4 deficient mice is believed to be prevention of the rapid inactivation of GLP-1 by DPP4 [14, 15, 34, 35]. Thus, the active form of circulating GLP-1 accumulates in DPP4 gko mice. In contrast, GLP-1 levels were comparable in FAPgko and WT mice on chow, or on HFD when fasting or following a glucose challenge (Fig. S11A). Therefore, the improved glucose homeostasis in FAPgko mice was not driven by preservation of active GLP-1.

Deletion of FAP did not result in compensatory regulation of related enzymes. Intrahepatic mRNA expression of *Dpp4*, *Dpp8* and *Dpp9* were unchanged compared to WT mice on chow or HFD, and were, as expected, abundant (Fig. S12). Intrahepatic *Dpp9* expression was significantly less in HFD compared to chow fed mice (Fig. S13). Furthermore, *Dpp9* expression strongly correlated with *Dpp4* and *Chrebp* expression in both WT and FAPgko liver (Fig. S13B, C).

### Differential intrahepatic expression of genes associated with lipogenesis and triglyceride and fatty acid uptake

*De novo* lipogenesis contributes to the pathophysiology of NAFLD as it contributes almost one third of the accumulated intrahepatic triglycerides in hepatosteatosis. Peroxisome proliferator-activated receptor gamma (Pparg) and glucokinase (*Gck*) are key genes involved in *de novo* lipogenesis [36, 37]. Intrahepatic lipogenic gene downregulation on HFD included *Gck* in FAPgko and FAPgki mice and *Ppar*g in FAPgko mice, compared to WT mice (Fig. 8A, B, F), implying that FAP is an upstream regulator of *de novo* lipogenesis.

**Fig. 8.**
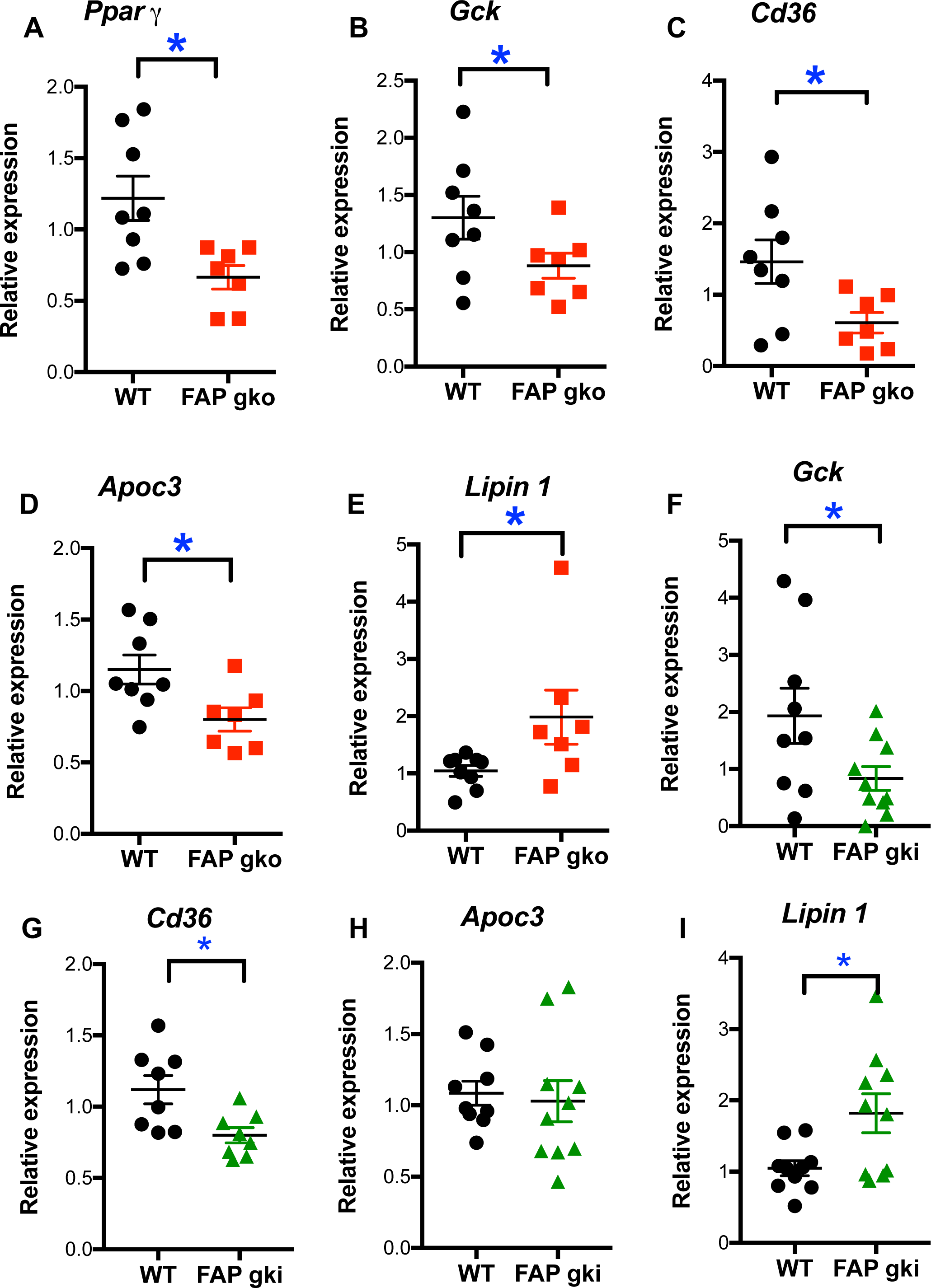
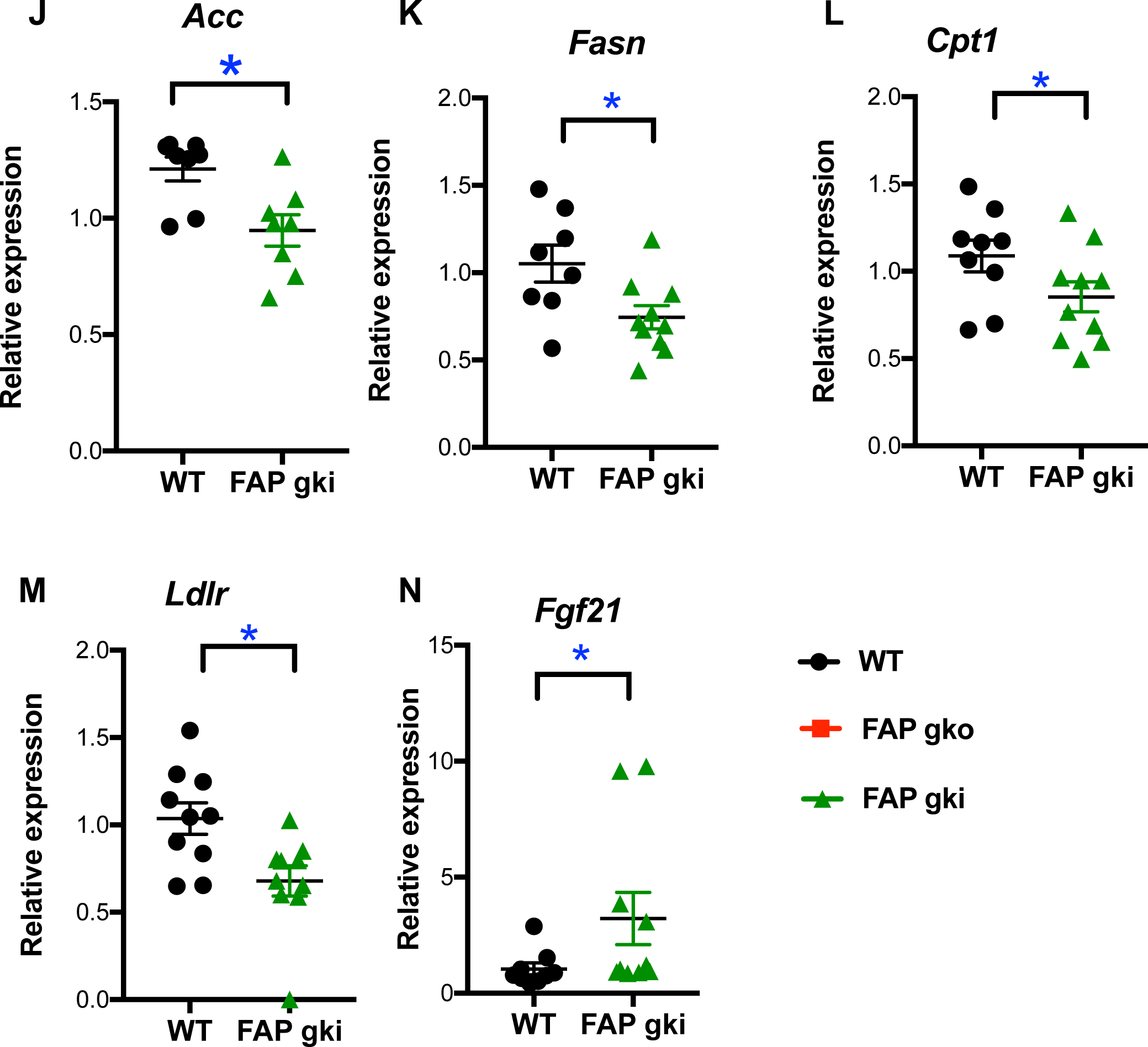
Differential intrahepatic mRNA expression of energy metabolism genes in FAP deficient DIO mice compared to WT mice. After 20 weeks of HFD mice were fasted overnight and most (A-F, H-M) were re-fed for 4 h. n=7-10 male (A-F, H-M) or 4-5 female (G) mice per group. Transcripts relative to housekeeper gene β-actin (A-F, H-M) or 18S (G). Individual replicates and mean ± SEM. *p<0.05.

*Cd36* and *Apoc3* are major components of triglyceride and fatty acid transport [38, 39]. On HFD, intrahepatic gene downregulation of *Cd36* occurred in FAPgko and FAPgki mice and of *Apoc3* in FAPgko mice, compared to WT HFD mice (Fig. 8C, D, G). Downregulation of *Ldlr*, *Acc, Fasn* and *Cpt1* in FAP deficiency (Fig. 8M) may also have contributed to lowered steatosis. CD36 immunopositivity in hepatocytes was less in FAPgki than WT liver (Fig. S14), concordant with the decrease in mRNA.

*Lipin-1* is a regulator of liver lipid metabolism, insulin sensitivity and glucose homeostasis [40, 41]. On HFD, intrahepatic *Lipin-1* expression was upregulated in FAPgko and FAPgki mice compared to WT mice (Fig. 8E,I), suggesting that the mechanism by which FAP deficiency decreased insulin resistance and liver lipid includes *Lipin-1*.

Decreased *Srebp1-c*(lipogenic gene) and increased *Ppar*a (lipolytic gene) are thought to be the key gene expression changes responsible for decreased lipogenesis in DPP4 gko mice [14]. We observed no change in *Srebp1-c* and *Ppar* a in FAPgko and FAPgki mice compared to WT mice on HFD (Fig. S11BE).

### Elevated FGF-21 in FAP deficient mice

The level of active FGF-21 increases in mice treated with an inhibitor of FAP enzyme activity [6], Therefore, *Fgf-21* upregulation (Fig. 8N) was possibly the most important differentially expressed gene, as it suggests that FAP deficiency upregulates *Fgf-21* expression in addition to suppressing FGF-21 degradation. The increased intrahepatic FGF-21 protein and its correlation with glucose AUC (Fig. 9A, B) may derive from both mechanisms, or gene transcription alone. Showing that FAP can prevent FGF-21 driven Erk phosphorylation in 3T3-L1 cells (Fig. 9C) further suggests that metabolic outcomes in FAP deficient mice are FGF-21 mediated. However, p-Erk levels were not altered in WAT (Fig. 9D), so there are probably additional mechanisms involved.

**Fig. 9.**
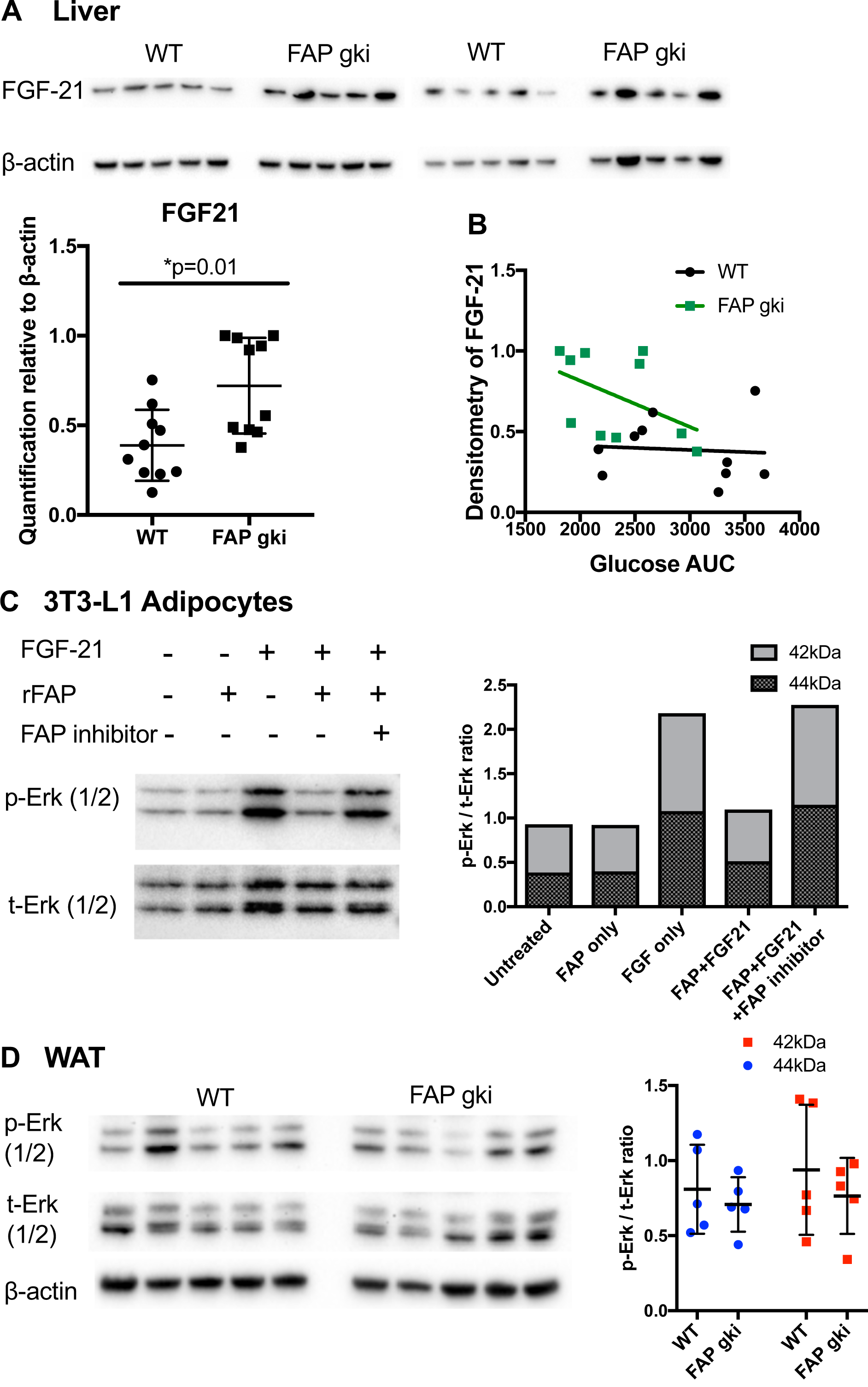
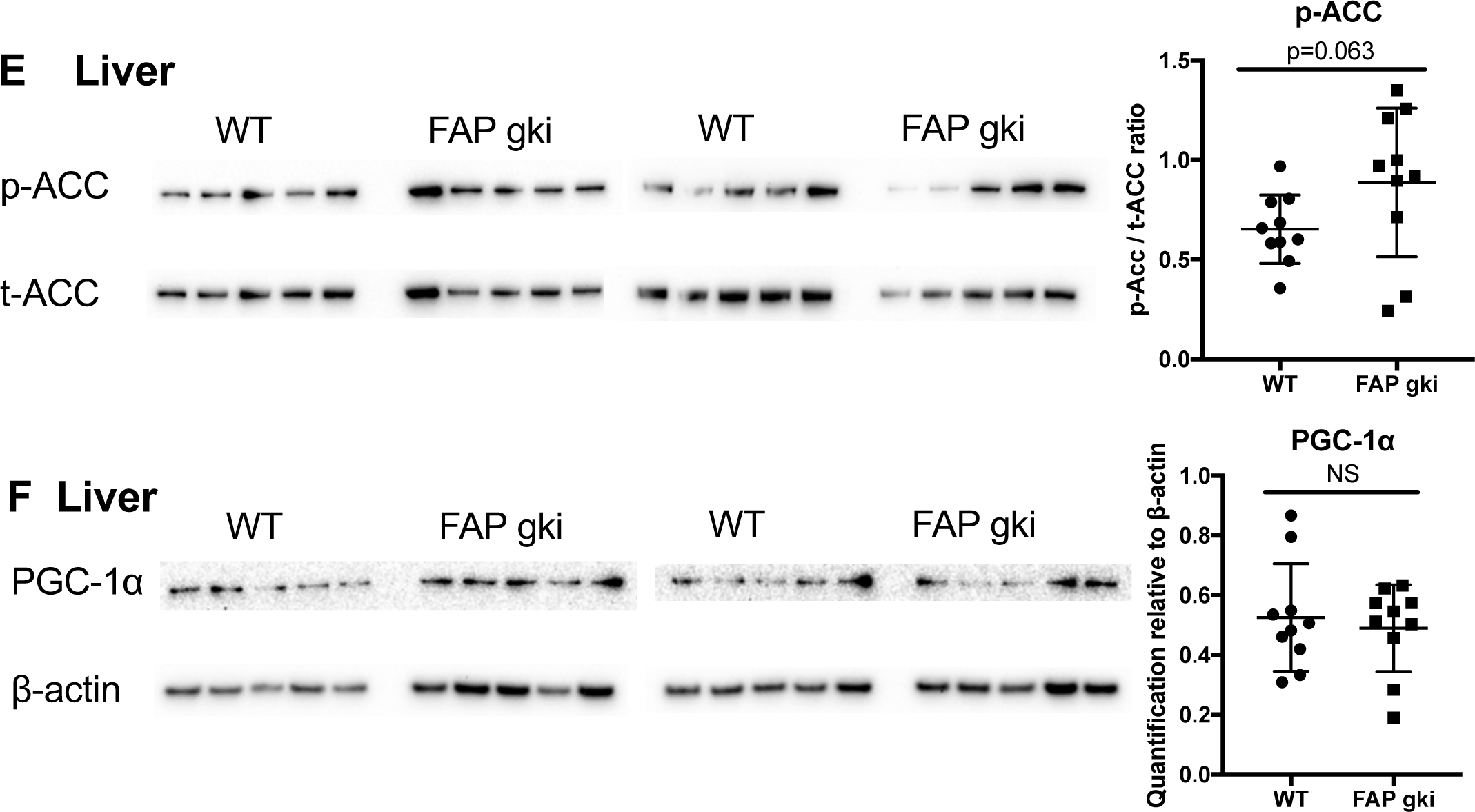
Increased hepatic FGF-21 expression in FAPgki mice and enhanced Erk signalling in 3T3-L1 cells. (A) Immunoblots of liver lysates from HFD fed WT and FAPgki mice showed increased FGF-21 expression in FAPgki mice (normalized to β-actin; n=10 per group). (B) Correlation of FGF-21 immunoblotting densitometry with GTT AUC. (C) 3T3-L1 cells that had been differentiated into adipocytes were incubated with full-length or FAP-cleaved FGF-21 at 500 ng/mL. FGF-21 was pre-incubated with PBS or recombinant FAP (substrate: enzyme ratio = 5:1) for 2 h, with or without FAP-selective inhibitor 3099 at 5 µM. Phosphorylated Erk (p-Erk) and total Erk (t-Erk) immunoblots and densitometry. Immunoblots are representative of two independent experiments. (D) Immunoblots of epididymal white adipose tissue (WAT) lysates from WT and FAPgki DIO mice showed increased Erk phosphorylation in FAPgki mice (n=5 per group). Densitometry data is p-Erk: tErk ratios. (E, F) ACC and PGC-1α immunoblots of liver lysates from WT and FAPgki DIO mice, with densitometry depicted as ratios of p-ACC: t-ACC (E) and PGC-1α:β-actin (F). N=10 per group.

### FAPgko mice had increased lipid metabolism

By indirect calorimetry, FAPgko and WT mice showed similar physical activity and energy expenditure on both chow and HFD (Fig. S15A-B). Therefore, the improved glucose and lipid metabolism in FAPgko mice was probably not associated with increased energy expenditure.

The RER represents the ratio of O_2_ consumption to CO_2_ production. FAPgko mice exhibited significantly less RER compared with WT mice, on either chow or HFD (Fig. 10A-D), which indicates a shift towards β-oxidation of lipid, rather than deriving energy from carbohydrate.

**Fig. 10.**
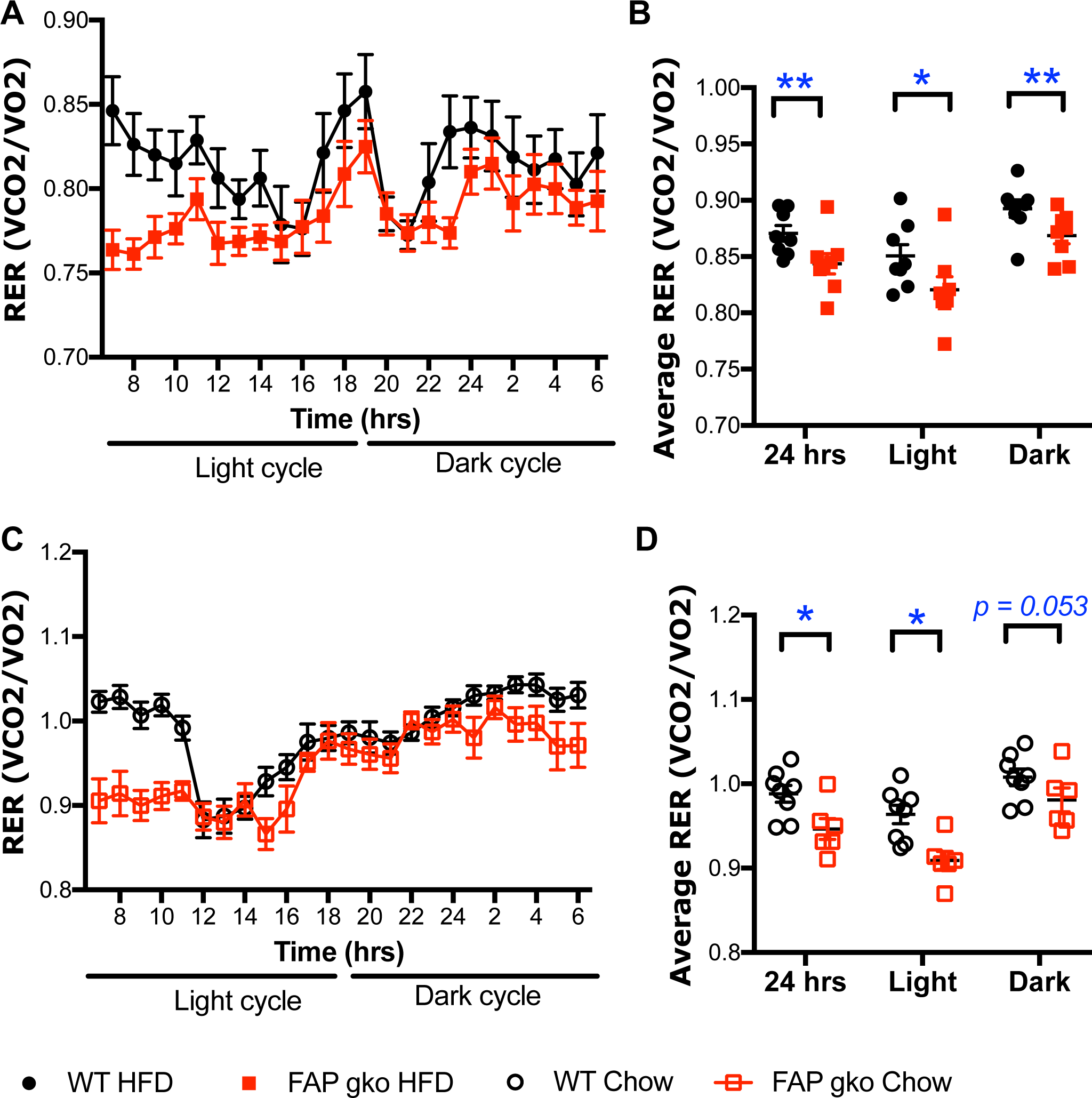
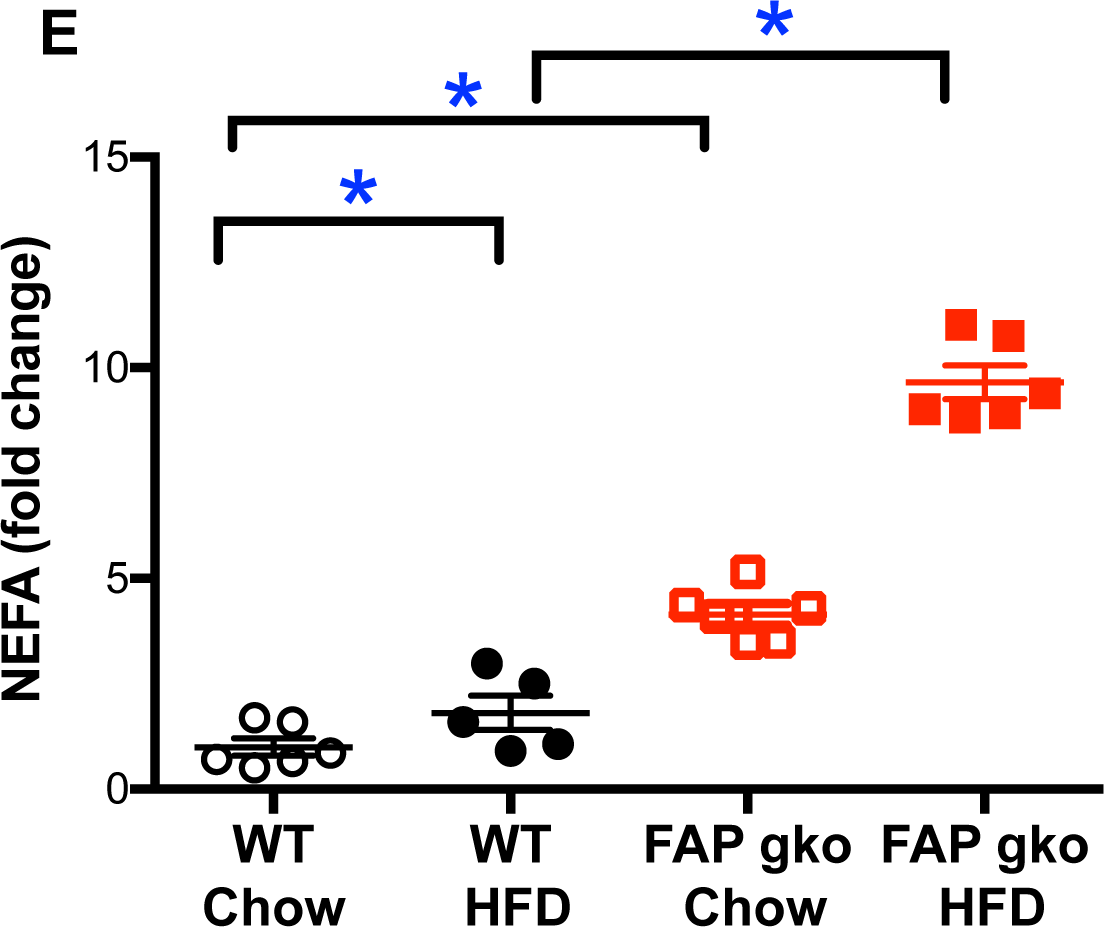
Respiratory exchange ratio and intrahepatic NEFA. Time course of respiratory exchange ratio (RER, an index of oxidative fuel) at 12 weeks of in-house HFD and 20 weeks chow (n=6-8 female mice per group) (A, C). Average RER from 24 hours, light phase (7am to 6pm) and dark phase (7pm to 6am) (B, D). Intrahepatic NEFA, as fold change from the WT chow average, was measured following 20 weeks of chow or HFD then overnight fasting (E). Mean ±SEM (A, C), or individual replicates and mean ±SEM (B, D, E). *p<0.05.

NEFA, also called free fatty acids, are generated by lipolysis, primarily in adipose tissue. Consistent with the RER data, FAPgko mice on either chow or HFD had elevated intrahepatic NEFA compared to WT mice on HFD (Fig. 10), which is concordant with increased intrahepatic lipid burning and less adipose tissue in FAPgko compared to WT mice.

Elevated fat oxidation is generally associated with elevated phosphorylation of ACC, which was observed in most of the FAPgki mice, and was not associated with PGC1a (Fig. 9E, F).

There was no difference in food intake between the two genotypes (data not shown). Faecal fat was measured as an indicator of fat absorption. No difference in faecal fat was detected in FAPgko compared to WT mice (WT = 72.93 ± 2.21 mg/g faeces; FAPgko = 70.27 ± 4.16 mg/g faeces), suggesting that FAP deficiency did not alter fat absorption.

### Elevated pancreatic FAP in human diabetes

Human pancreas samples from both type I and type II diabetes had elevated FAP enzyme activity compared to controls (Fig. 11A; Table S2.). FAP was colocalised to β-cells and not α-cells from human pancreas (Fig. 11B).

**Fig. 11.**
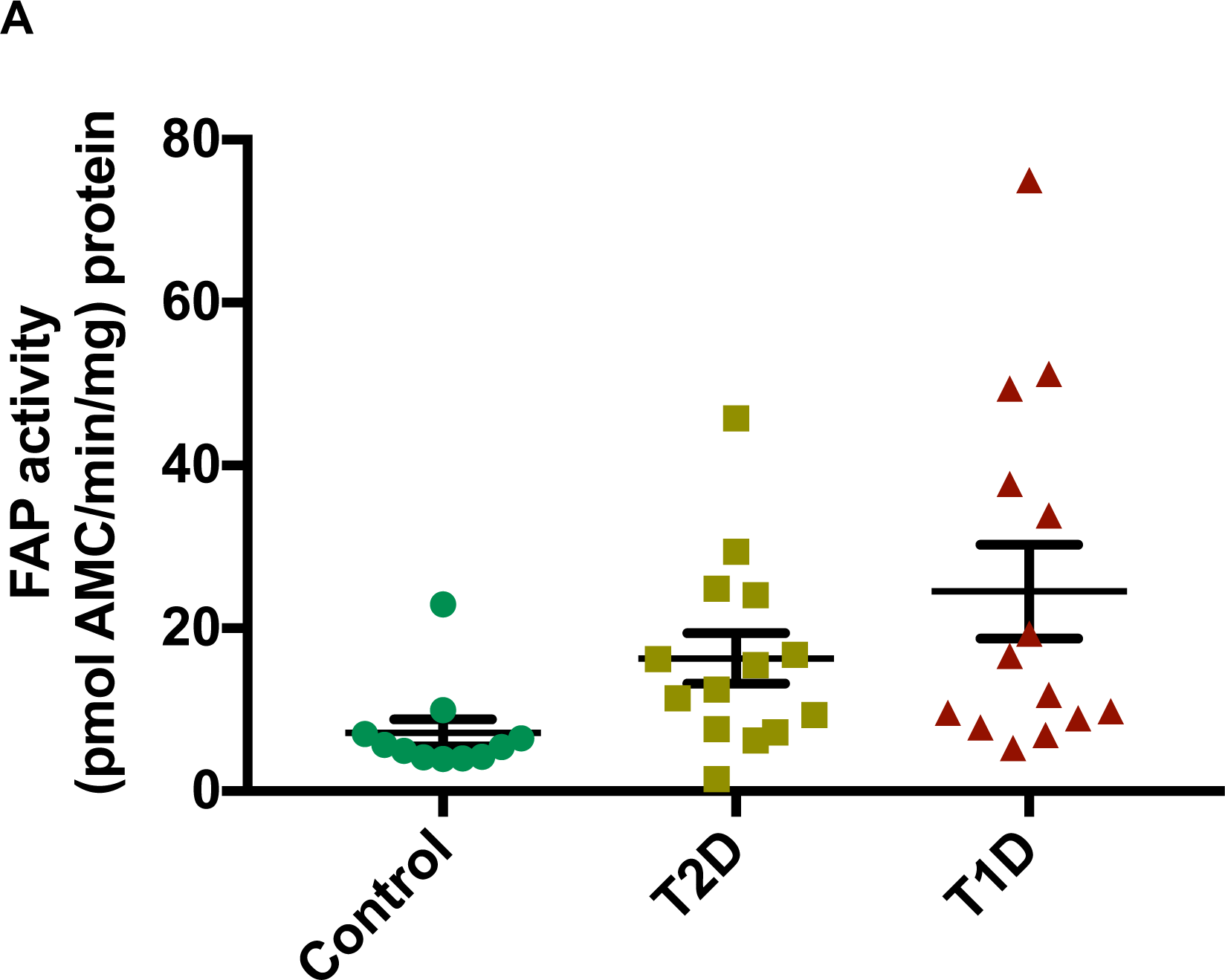
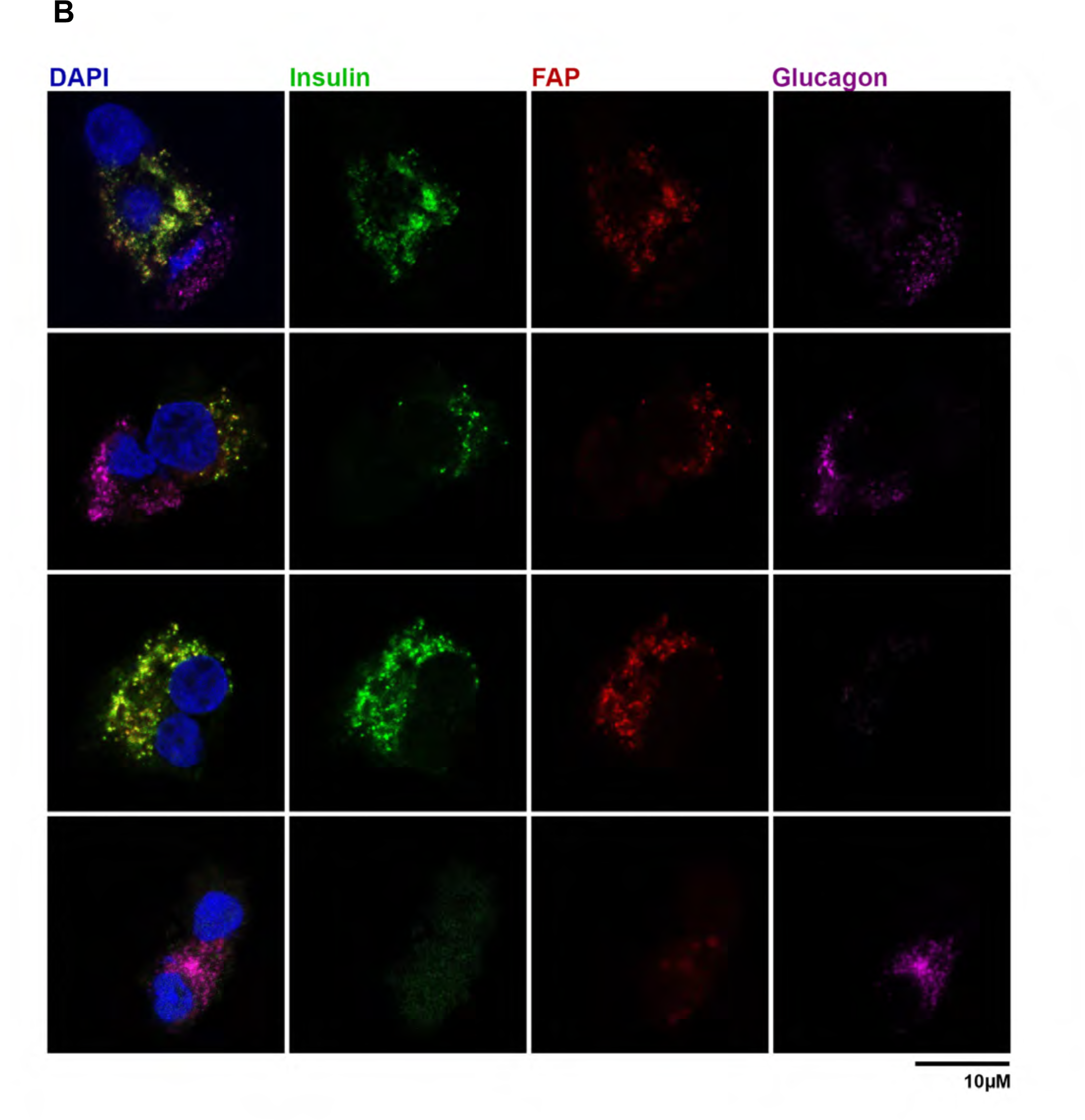
FAP in human pancreas: upregulation in diabetic pancreas and expression in pancreatic β-cells. FAP enzyme activity in pancreatic tissue extracts from individual controls and humans with type 1 or type 2 diabetes (A; mean ±SEM). Immunostaining for FAP (red), insulin (green; β-cells), glucagon (purple; α-cells) and DAPI (blue; nuclei) on human pancreatic cells on coverslips (B).

### Discussion

This study revealed a novel regulatory role of FAP in energy metabolism, and many aspects of this role. We showed for the first time that FAP is upregulated in pancreas from humans with diabetes and that FAP is present in pancreatic β-cells. We present the first demonstration that a specific FAP deficiency can reproducibly protect both male and female mice from obesity induced metabolic conditions. These include liver steatosis, hyperinsulinaemia, pancreatic β-cell hyperplasia, glucose intolerance, hypercholesterolaemia, visceral adiposity, elevated ALT and depressed adiponectin (Fig. S16). We further showed that loss of the enzymatic activity of FAP is sufficient to produce these outcomes, because a genetically modified mouse strain that lacks only the enzymatic activity of FAP recapitulated the phenotype of the global FAP knockout mice. The observed increases in intrahepatic lipid oxidation, FGF-21 and *Lipin-1* and *Fgf-21* expression in FAP deficiency appear to be key mechanisms by which FAP regulates energy metabolism. The evidence here indicates that FAP is an upstream regulator of lipogenesis and lipolysis and that FAP enzyme activity is required for hyperinsulinaemia in our DIO model. The findings were robust; derived from two FAP deficient mouse strains and several fat-laden diets.

FAP deficiency caused downregulated microvesicular steatosis, which is the more severe form of steatosis that is associated with liver injury, fibrosis progression and insulin resistance [42], so this was probably the cause of lowered ALT and AST. Macrovesicular steatosis rarely progresses to fibrosis [43]. Mitochondrial β-oxidation protects against microvesicular steatosis [43]. The reduced RER, increased intrahepatic NEFA, reduced microvesicular steatosis and elevated ACC and p-ACC point to increased β-oxidation in FAP deficient liver. Moreover, the reduced RER indicates a greater lipid to carbohydrate ratio as energy sources and increased intrahepatic NEFA and ACC are consistent with increased lipolysis. These data suggest that FAP negatively regulates β-oxidation, which would promote insulin resistance in obesity. Therefore, it is probable that improved β-oxidation in FAP deficient liver decreased the insulin resistance. The reduced sizes of WAT and BAT in FAP deficient mice are consistent with increased β-oxidation also occurring in adipose tissue. Showing that FAP regulates FGF-21 driven cytoplasmic signaling in 3T3-L1 cells points to FGF-21 as a mechanistic link between FAP and lipid regulation.

Our discovery that ablating the FAP enzyme activity is sufficient to generate these metabolic outcomes indicates that these outcomes are driven by bioactive natural substrates of FAP. The known metabolism-associated substrates of FAP are FGF-21, GLP-1, NPY and PYY [1, 3, 5, 20], and possibly nucleobindin-1 [44]. We showed here that GLP-1 activity is not altered and so is unlikely to be a physiological substrate of FAP. FGF-21 is a physiological substrate of FAP in primates [20] and mouse [6], and so FGF-21 is possibly the most relevant FAP substrate. We very significantly also found that FAP deficiency drives up *Fgf-21* expression, so FAP exerts two mechanisms of upregulating FGF-21. Mouse FGF-21 is poorly hydrolysed by FAP but FAP-mediated cleavage accelerates inactivation of murine FGF-21 [6, 20]. The changes in adiposity are small and food intake was unaltered, so appetite regulation by NPY, PYY or nucleobindin-1 are less likely mechanisms of FAP action.

The increased pancreatic FAP levels in human diabetes, the detection of FAP in pancreatic β-cells, and the limited β-cell hyperplasia in FAP deficient mice suggest that mechanisms by which FAP regulates glucose and insulin homeostasis may primarily occur in the pancreas. FGF-21 is a physiological FAP substrate [20] and its secretion is regulated by glucagon [45]. The hepatokine FGF-21 regulates lipid metabolism, increases intrahepatic mitochondrial β-oxidation and decreases liver steatosis [46]. Adiponectin, which was elevated in FAP deficient mice, can mediate FGF21 action [47]. Thus, FGF-21 may be a key component of the mechanism by which FAP regulates energy metabolism (Fig. S16).

The physiological and metabolic improvements in FAP deficient mice were associated with altered intrahepatic expression of lipid metabolism genes. Lipin-1 downregulates insulin resistance and FGF-21 increases β-oxidation. PPARg and glucokinase are lipogenic and CD36 and Apoc3 are essential for fatty acid and triglyceride uptake in the liver. These genes are upregulated in mouse models of obesity and liver steatosis and are implicated in human NAFLD [36, 37, 39, 48]. CD36 was downregulated at both protein and message levels, so CD36 regulation is probably downstream from FAP. Hence, less *de novo* lipogenesis, less triglyceride and fatty acid uptake and improved regulation of glucose and fatty acid metabolism could be responsible for less liver steatosis in FAP deficient livers.

DPP4 is the enzyme structurally most similar to FAP. DPP4 inhibition is a successful treatment for type 2 diabetes and is able to prevent liver steatosis and fibrosis [13, 18, 25, 49, 50]. The metabolic outcomes in the FAP deficient mice were similar to those in DPP4 gko and DPP4 inhibitor treated DIO mice [14, 15, 51]. Recently, a comparable phenotype has been observed in DIO mice treated with the non-selective FAP inhibitor Val-boro-Pro (Talabostat; PT100) [6]. Notably, the metabolic benefits of FAP downregulation are evident at 8 weeks or more of HFD and not chow in both the inhibitor – treated mice [6] and our genetically modified mice.

We found that FAP has mechanisms of action that are distinct from DPP4. DPP4 deficient mice in DIO have increased levels of active GLP-1, less food intake, increased energy expenditure, lower body weight, decreased *Srebp1-c* and increased *Ppar*a [14, 15, 17]. In contrast, these parameters were not involved in FAP deficient mice. Thus, FAP and DPP4 are associated with different pathways in energy homeostasis.

FAP has previously been found only in activated fibroblasts, so the new observation of its expression in β-cells commences a paradigm shift in understanding FAP functions. In DIO, FAP is probably acting on non-extracellular matrix substrates derived from β-cells and neighbouring cells, perhaps some that were identified in our recent degradomics study [1]. FAP is relatively abundant in mouse serum, so that is probably a major site of substrate degradation, including FGF-21. The dominant source of circulating FAP in healthy individuals, or in DIO, is not known.

Several genes involved in energy metabolism pathways, including fatty acid uptake, lipoprotein metabolism, adipokine transport and gluconeogenesis, are altered in DPP9 enzyme deficient neonate liver [52]. Val-boro-Pro inhibits both FAP and DPP9. In particular, intrahepatic *Lipin-1*, which downregulates insulin resistance, is differentially expressed in the *Dpp9* deficient mouse [52], fibrotic DPP4 deficient mice [25] and FAP deficient DIO mice (Fig. 9). These data together with our findings that intrahepatic DPP9 strongly correlated with *Dpp4* and *Chrebp* and that *Dpp9* expression was downregulated in DIO mice suggest that DPP4 gene family members FAP, DPP4 and DPP9 each have important, distinct roles in energy metabolism.

## Conclusion

In conclusion, this study revealed a novel function of FAP in regulating energy metabolism. We provide strong evidence that complete deficiency of FAP activity is beneficial and protected mice from DIO-induced impaired glucose homeostasis, hyperinsulinaemia and liver steatosis (Fig. S16). The evidence that FAP and DPP4 exert their functions through independent pathways imply that these enzymes may synergise in regulating metabolism. This novel role of FAP in metabolic disease together with the known benefit of DPP4 inhibition suggest that combined therapy targeting both FAP and DPP4 may offer potential for treatment of metabolic syndrome/diabetes.

## Acknowledgements

The authors thank the staff of the animal facility and the flow cytometry and imaging facility at Centenary Institute and the animal facility of Royal Prince Alfred Hospital for scientific and technical assistance. We thank Dr Andreas Schnapp and Dr Wolfgang Rettig for the FAPgko mouse strain. We thank staff of Ozgene, Professor David James and Dr Annabel Minard for advice and Dr Ben Crossett of Sydney Mass Spectrometry for advice and assistance.

## Author’s Contribution

S.C. performed majority of experiments, analysis, interpretation of data and manuscript drafting. S.S., X.W. provided concept and design and performed experiments. M.G.G., S.W., Y.L., K.E. performed some mouse experiments. H.E.Z., M.X. performed experiments and interpreted data. D.Y. contributed to design and generation genetically modified mice and concept and design of experiments. A.L., S.M., H.E.Z., N.T., J.G. critically revised the manuscript. M.K., B.Y. J.G., L.L., S.T., S.M., G.C., N.T., W.B. contributed to study design and interpretation. J.G., W.H., R.S. obtained and purified human islets. A.C., W.B. designed, performed and interpreted assays of human pancreas. G.C., N.T. performed and interpreted the indirect calorimetric studies. G.M., W.B. provided design, supervision and manuscript review. M.D.G. oversaw design, interpretation and writing and supervised the laboratory in which most studies took place.

## Abbreviations

ACC: acetyl Co-A carboxylase
*Apoc3*: apolipoprotein C3
AUC: area under curve
BAT: brown adipose tissue
*Cd36*: cluster of differentiation 36
DIO: diet induced obesity
DPP: dipeptidyl peptidase
FA: fatty acid
FAP: fibroblast activation protein-a
FGF-21: fibroblast growth factor-21
*Gck*: glucokinase
gki: gene knock in
gko: gene knock out
GLP-1: glucagon like peptide-1
GTT: glucose tolerance test
H&E: Haematoxylin and Eosin
HFD: high fat diet
HOMA: Homeostasis Model Assessment
ITT: insulin tolerance test
NAFLD: non-alcoholic fatty liver disease
NASH: non-alcoholic steatohepatitis
NEFA: non-esterified free fatty acid
NPY: neuropeptide Y
ORO: oil red O
*Ppar*g: peroxisome proliferator-activated receptor gamma
PYY: peptide YY
RER: respiratory exchange ratio
WAT: white adipose tissue
WT: wild type

## Lay Summary

High levels of insulin, cholesterol and sugar in the blood of people with fatty liver are major drivers of disease progression towards NASH. This study found that mice lacking the FAP enzyme are less prone to metabolic malfunctions including high blood sugar, insulin resistance and fatty liver when on a high-fat, high-sugar, high-cholesterol diet. Therefore, we predict that an appropriate FAP inhibitory compound could provide similar metabolic benefits.

